# Translational Downregulation of 5’ TOP mRNAs During T Cell Exhaustion

**DOI:** 10.64898/2025.12.15.694525

**Authors:** Erica S. Pledger, Navya Ambavaram, Lucas Ferguson, Jamie H. D. Cate

## Abstract

T cell exhaustion is a dysfunctional state that arises during chronic infections and cancer, characterized by impaired effector functions and sustained expression of inhibitory receptors. While transcriptional, epigenetic, and metabolic rewiring have been well documented in exhausted T cells, a comprehensive understanding of how translation is regulated in this state remains incomplete. To address this gap, we performed ribosome profiling and RNA sequencing on exhausted human CD8+ T cells to globally assess translational control. Our analyses reveal a marked repression of 5’ terminal oligopyrimidine (TOP) mRNAs during exhaustion. Unexpectedly, we demonstrate that this translational repression occurs despite evidence of elevated mTOR activity in exhausted T cells. These findings uncover a previously unknown layer of translational control in exhausted T cells.

## Introduction

CD8+ T cells, or cytotoxic T cells, play a critical part in immune surveillance by detecting and eliminating infected and malignant cells. Upon acute antigen stimulation, CD8+ T cells undergo rapid transcriptional, translational, and metabolic reprogramming to support clonal expansion and differentiation into effector cells (Araki et al., 2017; Kaech & Cui, 2012; Wolf et al., 2020). While the majority of these differentiated effector T cells eventually undergo apoptosis during the contraction phase of an immune response, a small surviving fraction gives rise to long-lived memory T cells. By contrast, CD8+ T cells encountering chronic antigen stimulation – such as in persistent viral infections or cancer – will gradually enter a state of dysfunction known as T cell exhaustion. This state is defined by the progressive loss of T cell effector functions, sustained expression of inhibitory receptors, and a failure to clear persistent antigen (Baessler & Vignali, 2024). Exhausted T cells fail to produce key cytokines, including IL-2, IFNγ, and TNFα, and upregulate inhibitory receptors that negatively regulate T cell activation, such as PD1, LAG3, and TIM3 (Baessler & Vignali, 2024). There has been significant progress in revealing the mechanistic underpinnings of T cell exhaustion, notably from an epigenetic, transcriptional, and metabolic perspective (Philip et al., 2017; Wu et al., 2023; Yao et al., 2019). However, an understanding of global changes in translation that occur during T cell exhaustion is largely lacking.

Protein synthesis is one of the most energy-intensive processes in a cell and is therefore subject to tight regulation. Translational control plays a critical role in CD8+ T cell function and fate decisions (Marchingo & Cantrell, 2022). Naive T cells are poised for activation by maintaining a reservoir of idle ribosomes and repressed mRNAs that can be rapidly translated following antigen stimulation (Wolf et al., 2020). Within just 6 hours after activation, T cells increase their protein synthesis rate by fivefold to support the burst in proliferation and increase in cell size (Wolf et al., 2020). This dramatic surge is driven by changes in the activity and expression of eukaryotic initiation factors (eIFs) as well as a substantial increase in ribosome biogenesis (Ahern et al., 1974; Howden et al., 2019; Mao et al., 1992; Miyamoto et al., 2005; Wolf et al., 2020). For example, naive T cells express little to no eukaryotic initiation factor 4E (eIF4E), but antigen stimulation induces its rapid expression (Howden et al., 2019; Mao et al., 1992). Protein synthesis remains high for several days until the peak of the effector response, at which point CD8+ T cells rapidly downregulate translation as they cease to divide (Araki et al., 2017).

A major regulator of translation is the serine/threonine kinase mTOR (mammalian target of rapamycin). mTOR regulates protein synthesis by phosphorylating translation inhibitors and ribosomal subunits (Marchingo & Cantrell, 2022). While this pathway globally promotes cap-dependent translation, a specific class of transcripts – 5’ terminal oligopyrimidine (TOP) mRNAs – are particularly sensitive to mTOR activity (Meyuhas & Kahan, 2015). TOP mRNAs contain a +1 C directly after the 5’ m^7^G cap followed by a series of 4 to 15 pyrimidines and mostly encode ribosomal proteins and other components of the translational machinery (Meyuhas & Kahan, 2015). In T cells, mTOR is a critical regulator of T cell differentiation, yet its influence on global protein synthesis appears more limited: pharmacological inhibition of mTOR during T cell activation only reduced protein mass by 26% in murine CD8+ T cells (Howden et al., 2019). Instead, mTOR signaling likely has a highly selective role in regulating the expression of TOP mRNAs. These transcripts are dynamically regulated during T cell differentiation – their translation is repressed in naive T cells, rapidly increased following antigen stimulation, and then strongly inhibited at the height of the effector response, right before the contraction phase (Araki et al., 2017; Stepanov et al., 2025; Wolf et al., 2020).

The state of protein synthesis in exhausted T cells has only recently begun to be explored. Studies in mice demonstrated that T cells undergoing exhaustion display a continuous rise in translational activity (Liu et al., 2025; Wang et al., 2025). Using ribosome profiling, Liu et al. found that hypertranslation of transcripts encoding oxidative phosphorylation genes contributed to mitochondrial dysfunction in exhausted T cells (Liu et al., 2025). Likewise, Wang et al. used mass spectrometry to reveal that heightened protein synthesis disrupts proteostasis, leading to the accumulation of stress granules and protein aggregates (Wang et al., 2025). These findings suggest that dysregulated translation is a key feature of exhausted T cells and underscore the need for further investigation into protein synthesis control during exhaustion.

Here, we chronically activated human CD8+ T cells *in vitro* to recapitulate the phenotypic signatures of T cell exhaustion. We then used ribosome profiling and RNA sequencing to probe the translational landscape of exhausted T cells. We observed a striking downregulation in the translation of TOP mRNAs when compared to effector T cells. Unexpectedly, we also provide evidence that mTOR is more active in exhausted T cells than in effector T cells. Collectively, our results reveal that exhausted CD8⁺ T cells possess a unique translational program, alongside the well-characterized epigenetic, transcriptional, and metabolic programs.

## Results

### *In vitro* chronically activated CD8+ T cells show hallmarks of T cell exhaustion

We first sought to establish a method for generating exhausted CD8+ T cells *in vitro*. We surveyed the literature for simple protocols that reproducibly produce large quantities of T cells that exhibit the molecular and phenotypic features of T cell exhaustion. An approach used by numerous groups is to chronically stimulate T cells with antibodies targeting the T cell receptor (TCR), bypassing the need for antigen stimulation (Belk et al., 2022; Canale et al., 2023; Lawton et al., 2024; Liu et al., 2025; Wang et al., 2025). Following this strategy, we purified CD8+ T cells from human peripheral blood mononuclear cells (PBMCs) of four healthy donors and activated them using anti-CD3 and anti-CD28 antibodies over the course of two weeks (Fig 1a). Our protocol employs a series of four 2-day activations separated by 2 days of rest, during which we remove the anti-CD3 and anti-CD28 antibodies to maintain cell viability and promote expansion (Fig 1a, chronic). For comparison, we also activated CD8+ T cells for 2 days and either let them rest for the remaining 12 days or re-activated them on the last 2 days to mimic naive and effector T cells, respectively (Fig 1a, acute and re-activated). These three conditions are herein referred to as T_chron_, T_re-act_, and T_acute_ cells.

**Figure 1.**
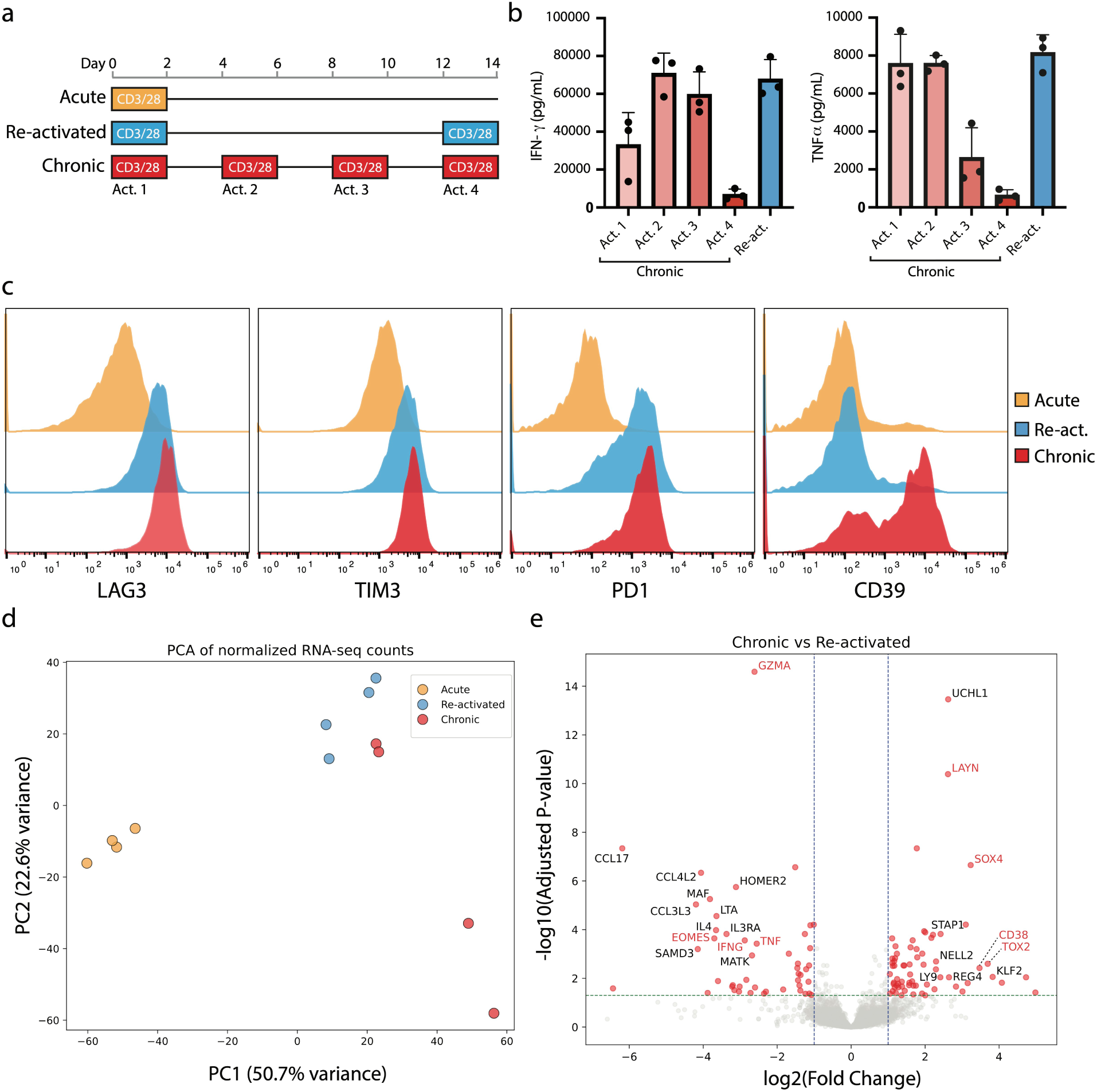
*In vitro* chronically activated CD8+ T cells show hallmarks of T cell exhaustion. **(a)** Schematic for *in vitro* generation of T_acute_, T_re-act_, and T_chron_ cells by anti-CD3 and anti-CD28 antibody activation. Each 2-day activation is indicated in T_chron_ (Act. 1, Act. 2, etc). **(b)** IFN-γ and TNFα secretion of T_chron_ and T_re-act_ measured by ELISA (n = 3 donors, error bars represent mean ± standard deviation). Of note, T_re-act_ (in blue) shows the final activation (ending on day 14), not the initial activation. **(c)** Representative flow cytometry histograms of the inhibitory receptors LAG3, TIM3, PD1, and CD39 (n = 4 donors). **(d)** Principal component analysis of RNA-seq data from T_acute_, T_re-act_, and T_chron_ cells in four donors. **(e)** Differential expression analysis of RNA-seq data comparing T_chron_ to T_re-act_ cells. Genes with significant changes in expression are in red (|log_2_(fold change)| > 1 and adjusted p-value < 0.05).

Next, we sought to determine if our T_chron_ cells exhibit signs of exhaustion, such as impaired cytokine production and expression of inhibitory receptors. We observed an almost complete loss of IFNγ and TNFα cytokine secretion by the fourth activation in T_chron_ as compared to T_re-act_ (Fig 1b). Notably, while the viability of T_chron_ modestly declined with each activation, the magnitude of cytokine loss was significantly greater than the loss of viability (Supplementary Fig 1a,b). We also observed high expression of the inhibitory receptors LAG3, TIM3, PD1, and CD39 in T_chron_ (Fig 1c). Of note, T_re-act_ also expressed high levels of these markers, which are known to be expressed transiently during T cell activation (Fig 1c). Interestingly, there was heterogeneity in expression dynamics amongst the inhibitory receptors in T_chron_, with some oscillating between high and low expression levels with each activation and rest interval (PD1), while the others gradually increased in expression throughout the course of the two weeks (LAG3, TIM3, and CD39) (Supplementary Fig 1c).

Exhausted CD8+ T cells possess a transcriptional program that is distinct from effector, naive, and memory T cells. We therefore performed RNA sequencing (RNA-Seq) on T_chron_, T_re-act_, and T_acute_ cells in four healthy donors to determine if T_chron_ cells exhibited canonical features of exhaustion. Principal component analysis (PCA) revealed that T_acute_ cells are transcriptionally distinct from T_re-act_ and T_chron_ cells (Fig 1d). T_re-act_ cells also form a well-defined cluster. Interestingly, in two donors, T_chron_ cells segregated from both T_acute_ and T_re-act_ cells, while in the other two donors they cluster near T_re-act_. This may be reflective of the heterogeneity within exhausted T cells. CD8+ T cells gradually differentiate from a more stem-like progenitor exhausted state to a terminally exhausted state. Thus, the two donors that cluster near the T_re-act_ cells may represent an earlier stage along this exhaustion trajectory compared to the other two donors. This observation also highlights the variability between donors and the distinct outcomes that can arise even when samples are subjected to the same chronic activation conditions.

Next, we identified differentially expressed genes by comparing T_chron_ to T_re-act_ cells (Fig 1e). In line with our observation that cytokine production is severely hindered in T_chron_ cells, both the *IFNG* and *TNF* transcripts were downregulated. We also observed upregulation of mRNAs encoding proteins previously linked to T cell exhaustion, such as the transcription factors TOX2 and SOX4, membrane glycoprotein LAYN, and ectoenzyme CD38 (DeRogatis et al., 2023; Good et al., 2021; Mahuron et al., 2020; H. Seo et al., 2019; Zheng et al., 2017). Interestingly, the mRNA for transcription factor Eomes (*EOMES*) was downregulated in T_chron_ cells in our data. Although T cell exhaustion is generally associated with elevated expression of Eomes, its expression varies across exhaustion states (W. Seo et al., 2021). Progenitor and intermediate exhausted T cells express lower levels of Eomes, whereas terminally exhausted cells exhibit high Eomes expression (Beltra et al., 2020). Thus, the downregulation of Eomes that we observe may be an indication that T_chron_ cells are closer to progenitor or intermediate exhausted T cells as opposed to terminally exhausted T cells.

### T_chron_ cells translationally downregulate TOP mRNAs

To investigate translation dynamics globally during T cell exhaustion, we also performed ribosome profiling in parallel to RNA-Seq on lysates from T_chron_, T_re-act_, and T_acute_ cells (Fig 2a). Ribosome profiling provides a snapshot of genome-wide ribosome occupancy on mRNAs by capturing and deep sequencing the ∼30 nucleotide ribosome-protected fragments (RPFs) left after nuclease digestion (Ingolia et al., 2009). The density of RPFs on a transcript is a proxy for the extent of protein synthesis (Brar & Weissman, 2015). Here, we used a simplified ribosome profiling workflow that uses P1 nuclease for the digestion step and ordered two-template relay (OTTR) for library preparation (Ferguson et al., 2023). These modifications to the original method greatly reduce the labor and input required to perform ribosome profiling, making it ideal for experiments where cell material is limited (Ferguson et al., 2023).

**Figure 2.**
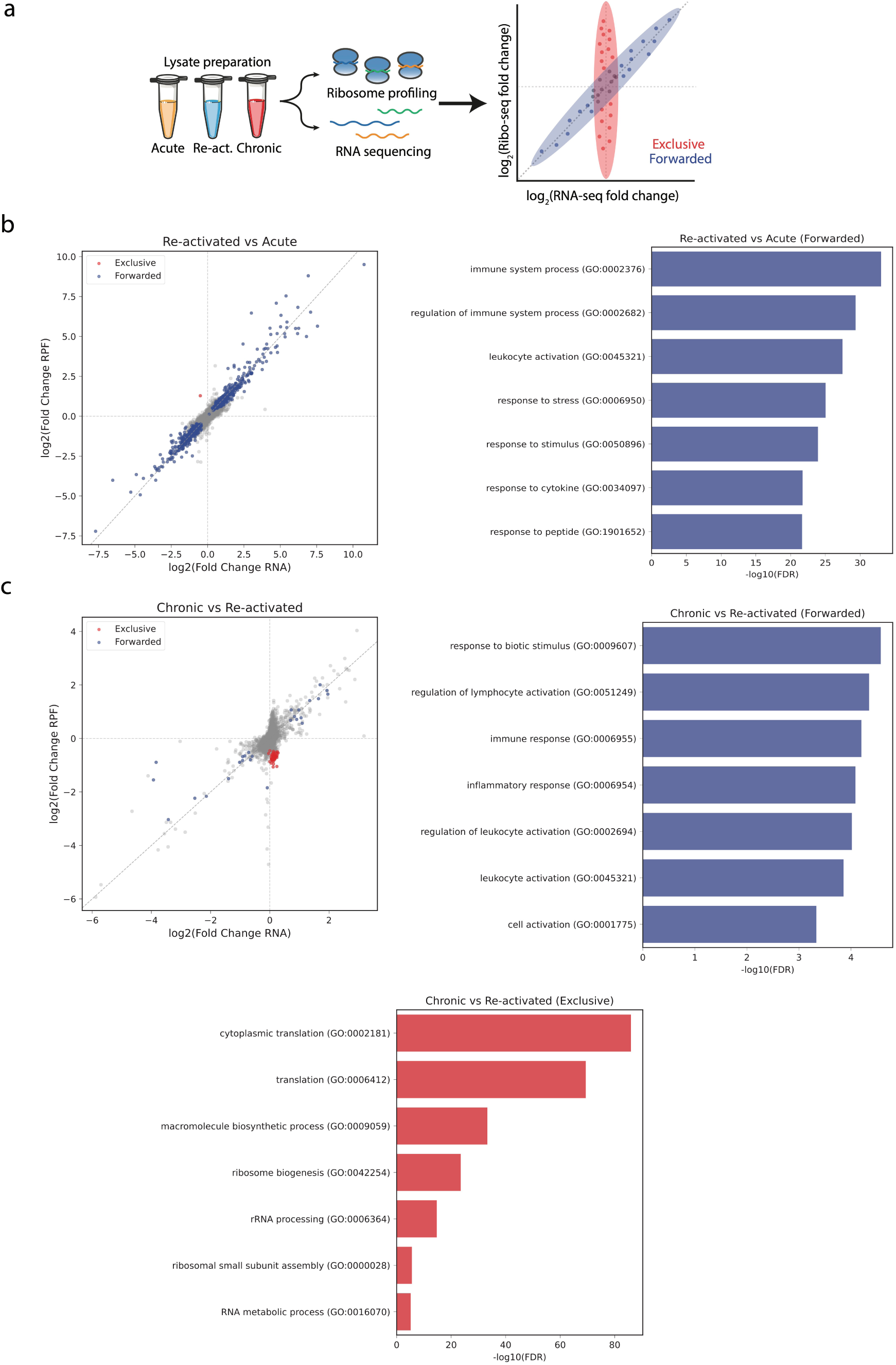
Transcriptional and translational regulation control distinct biological processes in exhausted T cells. (a) Schematic for studying translation regulation in T_acute_, T_re-act_, and T_chron_ cells. Lysates from each condition were prepared from four donors and RNA-Seq and ribosome profiling were performed in parallel. Differential expression analysis using deltaTE reveals genes that are forwarded (blue) and genes that are exclusive (red) (Chothani et al., 2019). Forwarded genes are regulated primarily at the transcript level, while exclusive genes are regulated primarily at the translational level. (b) Scatterplots of log_2_(fold change) RPF counts vs log_2_(fold change) RNA counts, comparing T_re-act_ to T_acute_ (n = 5851). Exclusive genes (n = 1) indicated in red and forwarded genes (n = 772) indicated in blue. Exclusive genes are defined as genes in which ΔmRNA is not significant (FDR > 0.05), ΔRPF is significant (FDR < 0.05), and ΔTE is significant (FDR < 0.05). Forwarded genes are defined as genes in which ΔmRNA is significant (FDR < 0.05), ΔRPF is significant (FDR < 0.05), and ΔTE is not significant (FDR > 0.05) (Chothani et al., 2019). To the right, GO analysis indicates the top biological processes associated with the forwarded genes. (c) Scatterplots of log_2_(fold change) RPF counts vs log_2_(fold change) RNA counts, comparing T_chron_ to T_re-act_ (n = 5851). Exclusive genes (n = 49) indicated in red and forwarded genes (n = 26) indicated in blue. GO analyses for the forwarded (blue) and exclusive (red) genes are also shown.

Our ribosome profiling results showed the expected read length distribution for eukaryotic RPFs, averaging around ∼35 nt (Supplementary Fig 2a) (Ferguson et al., 2023). The percentage of reads that mapped to mRNA (as opposed to rRNA, gDNA, etc) ranged from ∼10-30% (Supplementary Fig 2b). Principal component analysis of RPFs revealed a similar trend to the RNA-seq data – T_acute_ and T_re-act_ were distinct populations, while T_chron_ showed two donors that clustered with T_re-act_ while the other two donors formed a unique cluster (Supplementary Fig 2c and Fig 1d).

We then used our ribosome profiling and RNA-seq data to investigate gene regulation during T cell activation and T cell exhaustion programs (Supplementary Tables 1-3). To this end, we performed an integrative analysis of both our RNA-seq and ribosome profiling datasets using the deltaTE method to compare T_re-act_ to T_acute_ and T_chron_ to T_re-act_ (Chothani et al., 2019). Genes were sorted into two regulation classes: forwarded genes, where changes in RPF counts mirrored changes in mRNA counts and therefore indicate transcriptional regulation, and exclusive genes, where changes in RPF counts occur despite no change in mRNA counts and therefore indicate translational regulation (Chothani et al., 2019) (Fig 2a). Of note, there are two more regulation classes – buffered and intensified – which we did not observe in our analysis. When comparing T_re-act_ to T_acute_ cells, nearly all the differentially expressed genes (n = 773) fell under the forwarded category, suggesting that gene regulation occurs largely at the level of transcription during CD8+ T cell activation (Fig 2b and Supplementary Table 3). Consistent with this, gene ontology (GO) analysis revealed that the biological processes enriched among the forwarded genes were those related to immune system regulation and T cell activation (Fig 2b). By contrast, comparison of T_chron_ and T_re-act_ cells revealed only 26 forwarded genes, which were again enriched in processes related to immune function (Fig 2c and Supplementary Table 3). Interestingly, there were 49 genes categorized as exclusive that showed significant translational repression in T_chron_ relative to T_re-act_ cells (Fig 2c and Supplementary Table 3). GO analysis revealed that this group of translationally repressed genes is highly enriched for processes related to translation and ribosome biogenesis, including ribosomal protein genes (*RPS24*, *RPL32*), initiation factor genes (*EIF3E*), and translation elongation factor genes (*EEF1A1*, *EEF1B2*) (Fig 2c and Supplementary Table 3).

Next, we were interested in further investigating the group of exclusive genes that were translationally downregulated in T_chron_ cells. Importantly, the majority of the genes in this category are mRNAs that contain TOP motifs. We therefore hypothesized that all TOP mRNAs may be subject to dynamic translational control in T_chron_ cells. To test this, we analyzed the translation efficiencies (TE) of all genes using deltaTE (Chothani et al., 2019). TE is defined as the ratio of the number of RPFs mapping to the open reading frame (ORF) over the number of mRNA counts (Brar & Weissman, 2015; Chothani et al., 2019). We then compared TEs of T_chron_ to T_re-act_ cells and T_re-act_ to T_acute_ cells (Supplementary Fig 2d, Supplementary Tables 1-2). Strikingly, TOP mRNAs are significantly repressed relative to all other mRNAs when comparing T_chron_ to T_re-act_ cells (Fig 3a) (Philippe et al., 2020). In contrast, TOP mRNAs show no translation regulation during T cell activation (T_re-act_ vs T_acute_) (Fig 3a). Notably, the translational repression in T_chron_ occurs despite a mild increase in transcript abundance, indicating that downregulation is driven entirely at the level of translation (Fig 3b).

**Figure 3.**
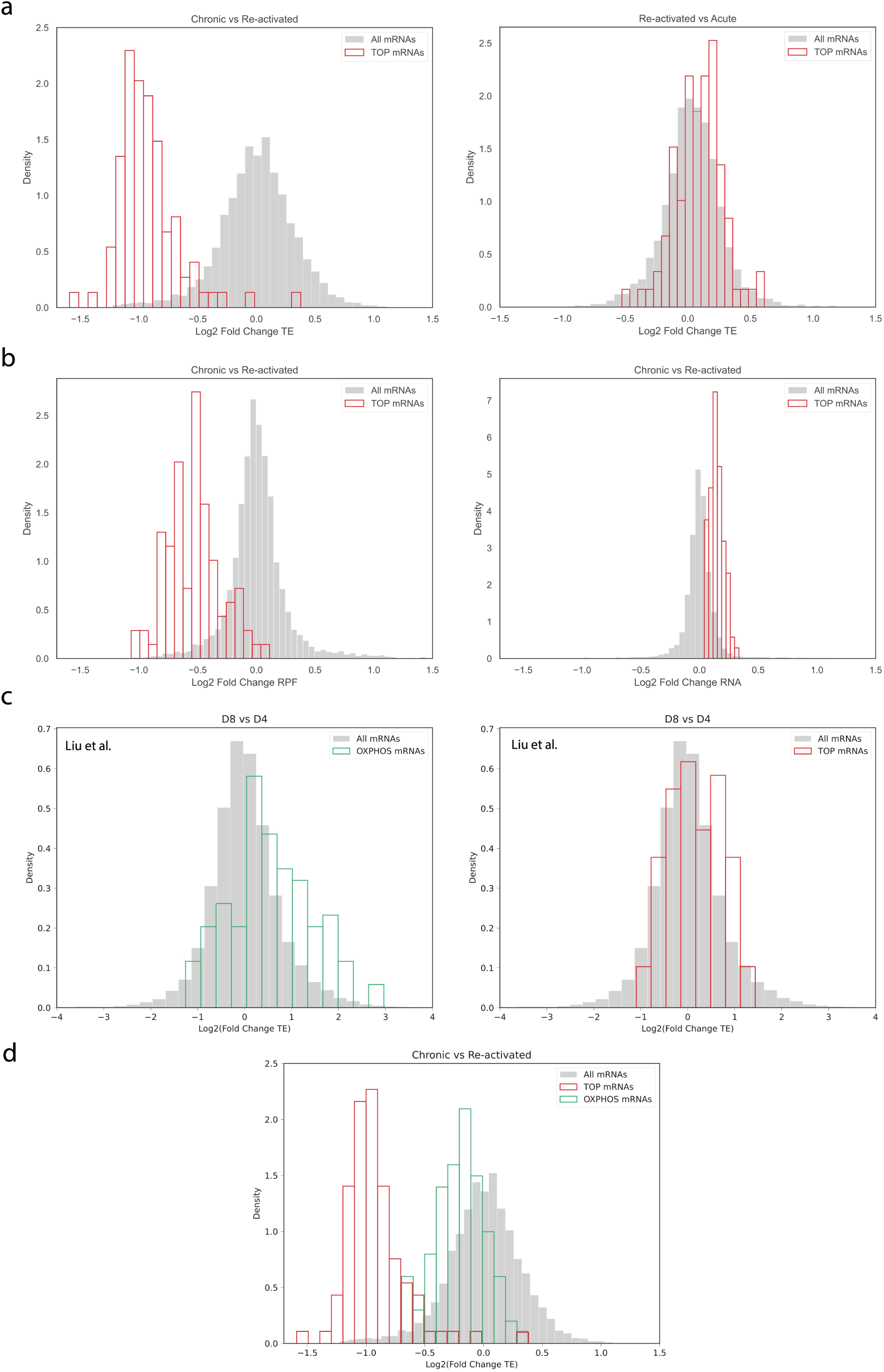
TOP mRNAs are translationally repressed in T_chron_ compared to T_re-act_ cells. **(a)** Histograms of log_2_(fold change) TE; all mRNAs in grey (n = 5851) and TOP mRNAs (n = 94) in red. Comparing T_chron_ to T_re-act_ on the left, comparing T_re-act_ to T_acute_ on the right. **(b)** Histograms of log_2_(fold change) mRNA and log_2_(fold change) RPF comparing T_chron_ to T_re-act_; all mRNAs in grey (n = 5851) and TOP mRNAs (n = 94) in red. **(c)** Histograms of log_2_(fold change) TE values comparing Day 8 (D8) to Day 4 (D4) chronically stimulated CD8+ murine T cells, taken from Liu et al., 2025; all mRNAs in grey (n = 9382), OXPHOS mRNAs in green (left, n = 139), and TOP mRNAs in red (middle, n = 94). **(d)** Histograms of log_2_(fold change) TE comparing T_chron_ to T_re-act_ from this study; all mRNAs in grey (n = 5851), OXPHOS mRNAs in green (n = 139), and TOP mRNAs in red (n = 94).

A recent study by Liu and colleagues used an *in vitro* chronic antigen stimulation model combined with a modified ribosome profiling approach to investigate translational dynamics during T cell exhaustion (Liu et al., 2025). We were therefore interested in assessing the similarities and differences observed in their results compared to those presented here. They observed hypertranslation of OXPHOS mRNAs driven by the RNA-binding protein LARP4 in their chronically stimulated CD8+ murine T cells (Fig 3c, left). However, they did not observe the translational repression of TOP mRNAs we observed (Fig 3c, right). In contrast to Liu et al., we did not observe the increase in translation efficiency of OXPHOS mRNAs in our T_chron_ cell data. In fact, we observed a mild repression in the translation of these mRNAs (Fig 3d). We explore possible reasons for this seeming discrepancy in the Discussion.

### mTOR activity is elevated in T_chron_

TOP mRNA translation is highly sensitive to the activity of mTOR. Given our observation that TOP mRNAs are translationally downregulated in T_chron_ cells, we sought to investigate the role of mTOR signaling during this process. To do so, we first assessed the state of mTOR activity using phosphorylation of ribosomal protein S6 (pRPS6) as a readout. We hypothesized that reduced mTOR activity in T_chron_ relative to T_re-act_ cells underlies the translational repression of TOP mRNAs that we observed in our ribosome profiling and RNA-seq data. Unexpectedly, we observed the opposite effect – pRPS6 levels were consistently higher in T_chron_ than in T_re-act_ cells throughout activations 2, 3, and 4 (Fig 4a and 4b). Notably, T_chron_ cells have greater levels of pRPS6 even after the last activation (activation 4), during which T_re-act_ cells were also subjected to the same activation (Fig 4a and 4b).

**Figure 4.**
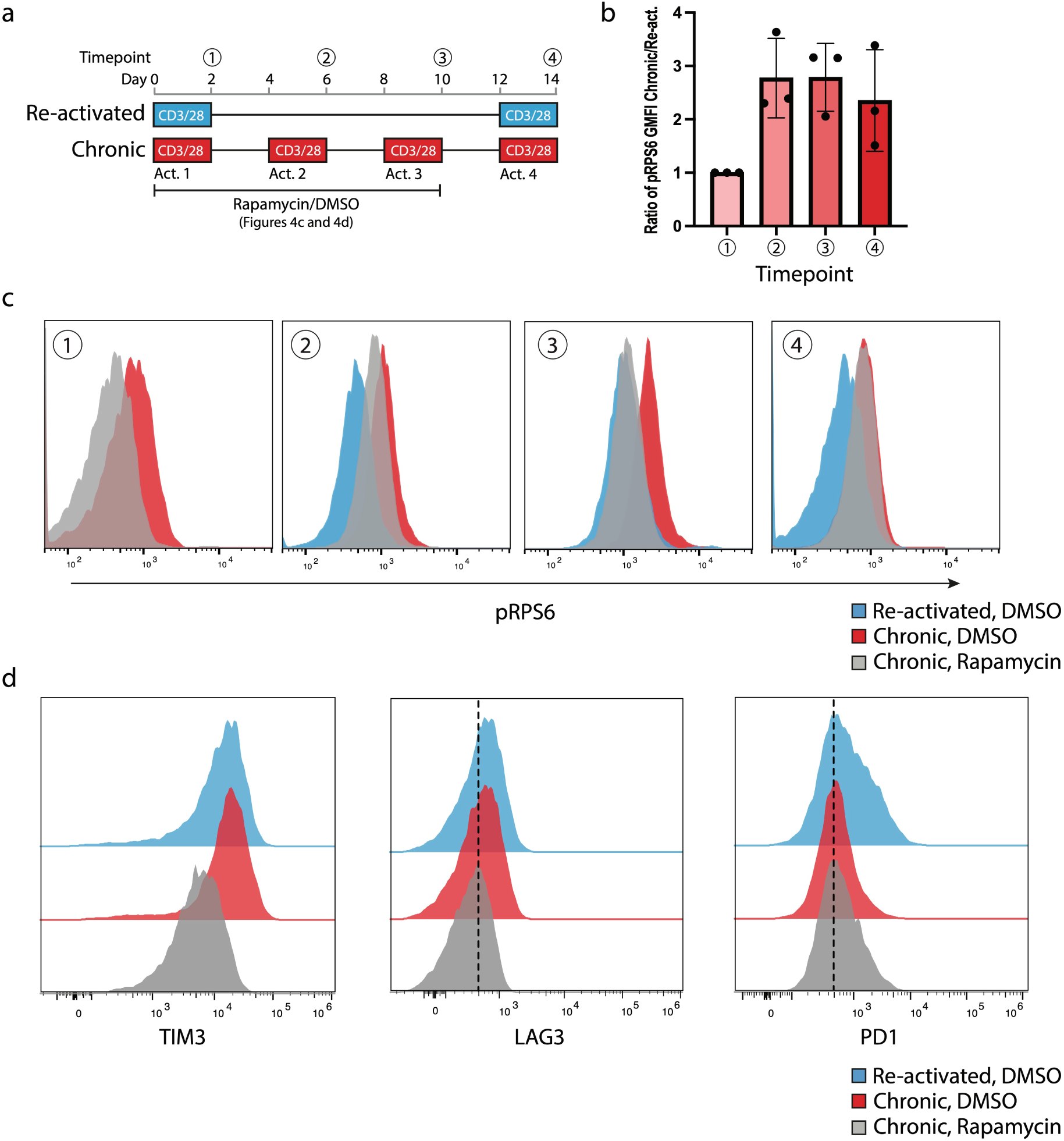
mTOR activity is elevated in T_chron_ compared to T_re-act_ cells. **(a)** Schematic for *in vitro* generation of exhausted T cells with timepoints at which flow cytometry measurements were taken, labeled (1-4). Cells were treated with rapamycin (or DMSO) during the first three activations for panels c and d. **(b)** The ratio of the geometric mean fluorescence intensity (GMFI) of phospho-RPS6 for T_chron_ over T_re-act_ at each timepoint (n = 3 donors, error bars represent mean ± standard deviation). **(c)** Representative flow cytometry histograms of phospho-RPS6 at each timepoint (n = 3 donors). **(d)** Representative flow cytometry histograms of PD1, LAG3, and TIM3 (n = 3 donors). The data shows the last timepoint (day 14).

Next, we used an inhibitor of mTOR, rapamycin, to test two alternative hypotheses based on our seemingly contradictory observations. First, given our observation that TOP mRNAs are repressed in T_chron_ cells, we hypothesized that inhibiting mTOR activity via rapamycin treatment would push these T cells towards an exhausted state by further repressing translation of TOP mRNAs. Alternatively, given our observation that mTOR activity is constitutively high in T_chron_ cells, we hypothesized that rapamycin treatment would put a brake on mTOR activity and slow down the progression of T cell exhaustion. We limited rapamycin treatment to the first three activations (Fig 4a), consistent with studies showing that rapamycin treatment improves outcomes when administered to mice during the initial phase of chronic infection, but has the opposite effect when administered after the establishment of exhaustion (Ando et al., 2023). Notably, rapamycin treatment reduced pRPS6 levels compared to the DMSO treated control in T_chron_ cells (Fig 4c), consistent with mTOR inhibition. However, after the last activation, at which point rapamycin had been removed from the media, pRPS6 levels in T_chron_ cells returned to the same level as the DMSO treated control T_chron_ cells (Fig 4c). We also observed a smaller cell size during rapamycin treatment of T_chron_ cells as measured by forward scatter (FSC), consistent with the role of mTOR in regulating cell growth (Supplementary Fig 3a, timepoint 2). Once again, this effect was transient and disappeared upon rapamycin removal (Supplementary Fig 3a, timepoint 4). Notably, rapamycin treatment improved cell viability throughout the two-week activation period as compared to T_chron_ cells treated with DMSO, as observed by periodic cell counting. We also used inhibitory receptor expression to assess the effect of rapamycin treatment on T_chron_ cells. Intriguingly, we observed a reduction in LAG3 and TIM3 expression, but not PD1 expression, in the rapamycin treated sample as compared to the DMSO treated control after the final activation (Fig 4d and Supplementary Fig 3b). This result better matches our second hypothesis – that rapamycin treatment effectively restrains mTOR activity to slow the progression of T cell exhaustion.

## Discussion

In this study, we performed RNA-sequencing and ribosome profiling in parallel to investigate changes in translation efficiency that occur during T cell exhaustion. Our *in vitro* method for generating exhausted CD8+ T cells effectively recapitulates key hallmarks of T cell exhaustion, including reduced cytokine production, inhibitory receptor expression, and changes in transcriptional programming. Moreover, similar results have been reported by other groups using chronic stimulation to model T cell exhaustion, highlighting the reproducibility and robustness of this method (Belk et al., 2022; Canale et al., 2023; Lawton et al., 2024; Liu et al., 2025; Wang et al., 2025). Importantly, this approach is simple and can produce the large number of cells necessary for genome-wide approaches such as ribosome profiling.

Our results reveal a striking downregulation in translation efficiency of genes encoding ribosomal proteins and other translation factors during T cell exhaustion. Intriguingly, the majority of the mRNAs encoded by these genes contain a unique feature at their 5’ end known as the TOP motif. Indeed, our data shows that nearly all the canonical TOP-motif-containing mRNAs were translationally repressed in T_chron_ cells compared to T_re-act_ cells. Notably, this repression occurred only at the translational level. In fact, these transcripts were mildly stabilized in the chronic condition as compared to the re-activated condition. Our findings provide new perspective on the dynamic regulation of TOP mRNAs in T cell biology. TOP mRNAs are some of the most abundant transcripts in naive T cells, and yet their translation is repressed until antigen stimulation, at which point their translation is rapidly induced to support ribosome biogenesis (Araki et al., 2017; Stepanov et al., 2025; Wolf et al., 2020). As the effector response reaches its peak, TOP mRNA translation is downregulated again as the cells stop proliferating (Araki et al., 2017). Our results suggest that a similar repression occurs during chronic T cell stimulation, perhaps indicating an adaptive response to help sustain cell survival under prolonged antigen exposure.

TOP mRNA translation is regulated by the mTOR pathway. Interestingly, we show that mTOR activity is sustained across all four activations in T_chron_ cells, seemingly in opposition to the inhibition of TOP mRNA translation that we observed. Several other groups have reported similar findings in mouse models of viral infection. During both acute and chronic infections, mTOR signaling initially peaks at the height of the effector response (Bengsch et al., 2016; Staron et al., 2014). However, in an acute infection, mTOR activity rapidly returns to baseline as the infection is cleared, whereas in a chronic infection, mTOR activity persists – albeit at a lower level – due to ongoing antigen stimulation (Bengsch et al., 2016; Staron et al., 2014). Notably, during chronic infection, T cells lose the ability to induce mTOR activity upon restimulation, while T cells responding to an acute infection maintain robust mTOR signaling capabilities (Bengsch et al., 2016; Staron et al., 2014). Even within exhausted T cell populations, variation in mTOR activity leads to different functional outcomes. Progenitor exhausted T cells exhibit suppressed mTOR activity in the earlier stages of the T cell response compared to their more terminally exhausted counterparts, which preserves their long-term fitness (Gabriel et al., 2021). Taken together, these findings highlight the dynamic role mTOR plays in governing T cell exhaustion and underscore the need for further investigation into how TOP mRNA regulation intersects with mTOR signaling.

Recent insights into the mechanism of TOP mRNA inhibition help explain how TOP mRNAs can be repressed even under conditions of high mTOR signaling. A key player in this mechanism is La-related protein 1 (LARP1), an RNA-binding protein that has been identified as the mTOR-sensitive repressor of TOP mRNA translation (Fonseca et al., 2018). Under conditions of mTOR inhibition, LARP1 binds the 5’ m^7^G cap and adjacent TOP motif of TOP mRNAs, blocking assembly of the eIF4F complex and thereby inhibiting translation initiation (Lahr et al., 2017). Initial models suggested that mTOR regulates LARP1’s binding activity by phosphorylating it at multiple residues and causing its disassociation from TOP mRNAs (Hong et al., 2017). However, more recently it was found that LARP1-TOP complexes also associate with 80S ribosomes to form repressed TOP-80S complexes under mTOR inhibition (Saba et al., 2024).

Intriguingly, Saba et al. showed that TOP-80S complexes also form during any condition which leads to greater numbers of free, idling ribosomes. This could occur even under conditions of normal mTOR activity, for example, by knocking down eIF4E to inhibit translation initiation. It is therefore possible that LARP1 may be capable of inhibiting TOP mRNA translation during T cell exhaustion even under conditions of high mTOR activity by “sensing” the presence of free ribosomes. This is a particularly attractive model in conjunction with evidence from Wolf et al. that naive T cells contain both large numbers of idling ribosomes and repressed mRNAs, the majority of which are regulated by mTOR. Further research will be necessary to determine if exhausted T cells also possess an abundance of free ribosomes.

LARP1’s role in regulating TOP mRNA expression is two-pronged – in addition to repressing their translation, LARP1 also stabilizes these transcripts in a highly selective manner (Gentilella et al., 2017). This was evident in our data, in which TOP mRNA TE was lower in T_chron_, despite transcript levels being slightly increased. LARP1 may also be essential in simultaneously stabilizing and repressing TOP mRNAs in naive T cells (Wolf et al., 2020). This also aligns with a previous study that reported transcriptional upregulation of ribosomal proteins and translation machinery during T cell exhaustion, and highlights that transcript levels may not accurately reflect protein expression (Yao et al., 2019).

We further probed the connection between TOP mRNAs, mTOR, and T cell exhaustion by using rapamycin to inhibit mTOR activity. Our results show that rapamycin treatment slows the progression of exhaustion, as evidenced by a reduction in inhibitory receptor expression. Similar observations have been reported by other groups. Rapamycin treatment during the early stages of a mouse model for chronic viral infection leads to better outcomes by promoting the formation of stem-like CD8+ T cells (Ando et al., 2023; Gabriel et al., 2021). In contrast, inhibiting mTOR activity later in the infection, when exhaustion was already established, repressed immunity by decreasing the numbers of effector-like T cells (Ando et al., 2023; Gabriel et al., 2021). Further work will be necessary to determine how mTOR inhibition affects TOP mRNA regulation during the exhaustion program.

We were also interested in comparing our ribosome profiling results to those published recently by Liu et al. (Liu et al., 2025). Their findings contrasted to ours. While they reported hypertranslation of OXPHOS and cytoplasmic translation genes during chronic activation, we observed the inverse – a mild repression of OXPHOS mRNAs and a marked repression of TOP mRNAs. These discrepancies could be due to differences in experimental design. Liu et al. activated murine CD8+ T cells with anti-CD3 antibodies continuously for 8 days without rest, whereas our study used human CD8+ T cells activated with anti-CD3 and anti-CD28 antibodies for 14 days with intermittent rest periods. These differences in experimental design complicate direct comparisons. Nonetheless, the stark contrast is intriguing and could reflect interesting biology. A possible explanation is that Liu et al. examined an earlier stage in T cell exhaustion and therefore captured a snapshot of the translational landscape that precedes what we observed. At this earlier stage, hypertranslation of translation machinery drives excess ribosome accumulation. In the later stages of exhaustion, the large number of idling ribosomes is sensed by LARP1, which causes formation of repressive TOP-80S complexes and consequent translational inhibition of TOP mRNAs. A future study covering a more comprehensive time interval – from the onset of exhaustion until the late stages – will be required to reconcile the Liu et al. results with the results reported here.

Finally, we synthesize the findings of several groups, as well as those presented here, to present a model for the role of translation regulation of TOP mRNAs during CD8+ T cell differentiation and exhaustion (Fig 5a). TOP mRNAs are abundant but repressed in naive T cells (Stepanov et al., 2025; Wolf et al., 2020). Naive T cells also contain large numbers of idle ribosomes, a strategy that enables a rapid transition to a translationally active state upon TCR stimulation (Wolf et al., 2020). Following antigen stimulation, TOP mRNAs are rapidly recruited to polysomes and translated in an mTOR-dependent manner as the cells clonally expand and differentiate into effector T cells (Araki et al., 2017; Wolf et al., 2020). Araki and colleagues demonstrated that CD8+ T cells downregulate TOP mRNAs at the peak of the effector response following antigen clearance (Araki et al., 2017). Notably, even among CD8+ effector T cells, terminal effectors – those that will eventually undergo apoptosis during the contraction phase – and memory precursors – those that will differentiate into long-lived memory T cells – exhibited distinct translational regulation of TOP mRNAs (Araki et al., 2017). In contrast, during chronic antigen exposure, TOP mRNAs remain highly translated (Araki et al., 2017; Liu et al., 2025; Wang et al., 2025). We hypothesize that sustained hypertranslation at this stage eventually triggers a feedback mechanism leading to TOP mRNA repression during the later stages of exhaustion, even in the presence of high mTOR activity.

**Figure 5.**
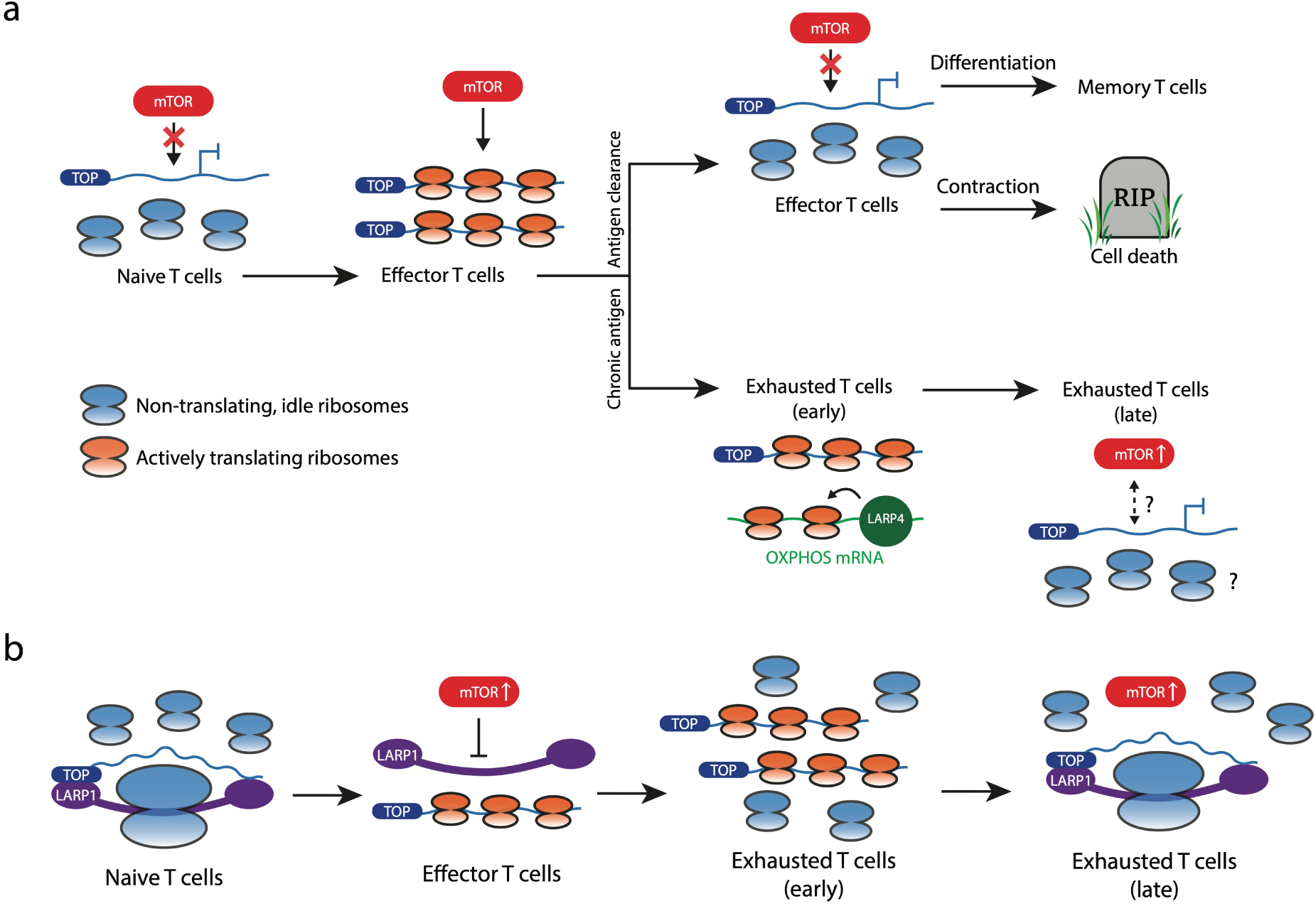
Model of translation regulation during T cell exhaustion. **(a)** Schematic of TOP mRNA regulation during the stages of T cell differentiation. Naive T cells contain large numbers of free ribosomes, and TOP mRNAs are repressed (Wolf et al., 2020). Upon antigen stimulation, TOP mRNAs are rapidly engaged by ribosomes and translated within polysomes (Araki et al., 2017; Stepanov et al., 2025; Wolf et al., 2020). During an acute infection, TOP mRNAs are repressed following antigen clearance, coinciding with the peak of the effector response – a process sensitive to mTOR activity (Araki et al., 2017). During the earlier stages of a chronic infection, TOP mRNAs and OXPHOS mRNAs are highly translated (Araki et al., 2017; Liu et al., 2025; Wolf et al., 2020). The hypertranslation of OXPHOS mRNAs is mediated by the RNA-binding protein LARP4 (Liu et al., 2025). In contrast, at later stages, TOP mRNAs are repressed despite high mTOR activity. **(b)** Hypothetical model of TOP mRNA regulation by LARP1 during T cell exhaustion. In naive T cells, LARP1 forms TOP-80S complexes that stabilize TOP mRNAs while repressing their translation. Upon antigen stimulation, LARP1’s repression is relieved by mTOR, allowing efficient translation of TOP mRNAs. During chronic antigen stimulation, hypertranslation of TOP mRNAs leads to an excess of free ribosomes. At later stages, LARP1 senses the presence of free ribosomes and forms TOP-80S complexes to repress TOP mRNAs, despite high mTOR activity.

We also propose a role for LARP1 in regulating TOP mRNA translation throughout T cell differentiation (Fig 5b). In naive T cells, where TOP mRNAs are translationally repressed and large pools of idle ribosomes are present, we hypothesize that LARP1-mediated TOP-80S complexes help maintain these cells in a poised, activation-ready state. In effector cells, the induction of mTOR activity following antigen stimulation relieves repression of TOP mRNAs by LARP1, which allows for their rapid translation. Under conditions of chronic stimulation, dysregulated TOP mRNA translation drives excess ribosome production. The resulting buildup of idle ribosomes is detected by LARP1, which triggers formation of repressive TOP-80S complexes, thus resulting in the translational repression of TOP mRNAs observed in this study. These hypotheses raise several compelling avenues for future work in understanding the mechanisms behind translation regulation in naive, effector, and exhausted T cells.

## Methods

### Cell culture

Human primary T cells were maintained in ImmunoCult™-XF T cell expansion medium (STEMCELL Technologies). Recombinant human IL-2 (Peprotech) was added to the media at the indicated concentrations. Cells were maintained between 2 x 10^5^ – 1 x 10^6^ cells/mL.

### Isolation of human primary T cells

Peripheral blood mononuclear cells (PBMCs) were isolated from human peripheral blood leukopaks (STEMCELL Technologies). Briefly, the contents of the leukopak were combined with an equal volume of MACS buffer (PBS, 2% FBS, 1 mM EDTA) and spun down (15 min, 100 xg, Dec = 1), following which the pellet was washed 1-2 times (5 min, 400 xg) in the MACS buffer. Bulk (CD3⁺) T cells or CD8⁺ T cells were isolated from the PBMCs by magnetic negative selection using the EasySep™ Human T cell isolation kit or Human CD8⁺ T cell isolation kit (STEMCELL Technologies), respectively, following the manufacturer’s protocol. For future use, cells were frozen in CryoStor® freezing medium (BioLife Solutions) and treated in the same manner as freshly isolated cells once thawed.

### In vitro chronic activation of CD8+ T cells

To model T cell exhaustion *in vitro*, CD8⁺ T cells were cultured under repeated stimulation conditions, as shown in Figure 1a. Cells were stimulated using plate-bound anti-CD3 (clone UCHT1) (Tonbo Biosciences (now Cytek)) at 5 µg/mL and soluble anti-CD28 (clone CD28.2) (Tonbo Biosciences (now Cytek)) at 2.5 μg/mL antibodies for 48 hours in ImmunoCult™ T cell expansion medium (STEMCELL Technologies) supplemented with recombinant human IL-2 (200 U/mL). After stimulation, cells were centrifuged and resuspended in fresh medium containing IL-2 (200 U/mL) and expanded for 48 hours. This stimulation and rest cycle was repeated four times over the course of two weeks. Cell lysates were collected 48 hours after the final stimulation. This condition is referred to as the “chronic” stimulation group and models T cell exhaustion.

In parallel, two additional conditions were maintained. The “acute” condition received only a single 48-hour stimulation at the beginning of the culture period, followed by expansion in IL-2-containing medium without further stimulation. The “re-activated” condition received an initial stimulation followed by expansion, and then a final stimulation at the end of the two-week period. All cultures were maintained under identical IL-2 concentrations (200 U/mL) and medium conditions throughout.

### Rapamycin treatment

Rapamycin was added to the media during both the stimulation and rest stages at 10 nM during the first three activations in T_chron_. As a control, we included a condition in which an equal volume of DMSO as in the rapamycin treatment was added to the media during the first three activations.

### Enzyme-linked immunosorbent assay (ELISA)

We used ELISAs to quantify cytokine production (Interferon gamma (IFN-γ) and Tumor Necrosis Factor alpha (TNF-α)) by T_chron_ and T_re-act_ following each stimulation. At each stimulation, cells were counted and plated in triplicate at 2 x 10^5^ cells/well in a 96-well flat bottom plate coated with anti-CD3 antibody (5 µg/mL). Anti-CD28 antibody (2.5 µg/mL) and IL2 (200 U/mL) were added directly to the media containing the cells. After 48 hours, the cell supernatants from these wells were collected and stored in the -80°C freezer until use. Cell supernatants were diluted appropriately and the ELISA MAX™ Human IFN-γ and TNF-α Sets (BioLegend) were used to determine IFN-γ and TNF-α concentrations following the manufacturer’s protocol. Absorption at 450 nm was measured using a TECAN Spark plate reader. The IFN-γ and TNF-α standards provided in the kits were used to generate a standard curve. GraphPad Prism was used to fit the standard curve.

### Flow cytometry

Flow cytometric analysis was performed on an Attune NxT Acoustic Focusing Cytometer (ThermoFisher). Briefly, cells were pelleted in a 96-well round bottom plate and washed with PBS before staining. For intracellular staining, cells were fixed and permeabilized using the Cytofix/Cytoperm Fixation/Permeabilization kit (BD Biosciences) according to the manufacturer’s instructions. Cells were stained with the following antibodies and live/dead stain diluted in PBS: LIVE/DEAD fixable near-IR (Invitrogen), CD8 (BioLegend, clone SK1), PD-1 (BioLegend, clone EH12.2H7), TIM3 (BioLegend, clone A18087E), LAG3 (BioLegend, clone 11C3C65), CD39 (BioLegend, clone A1), and RPS6 phospho Ser235/236 (BioLegend, clone A17020B). Cells were stained for surface markers on ice for 20 minutes in the dark. Cells were washed twice in PBS and resuspended in 150 µl of PBS for analysis. Flow cytometry data was analyzed using FlowJo software.

### RNA-seq

Total RNA was extracted from the same lysates collected for ribosome profiling using the Direct-zol RNA miniprep kit (Zymo Research). Library preparation and sequencing were carried out by Novogene using an Illumina NovaSeq X Plus 150PE platform. The FASTQ files provided by Novogene were already trimmed and filtered. Reads were aligned to a custom transcriptome that includes both the NCBI RefSeq MANE v0.95 transcript set and the 13 mitochondrial mRNA sequences from Ensembl (Ferguson et al., 2023). We used Salmon to quantify transcript-level abundances (Patro et al., 2017). PyDESeq2 was used for differential expression analysis (Muzellec et al., 2023).

### Ribosome profiling

Samples were prepared according to the protocol described in Ferguson et al. and on the Karnateq website (Ferguson et al., 2023). Briefly, on the day of collection, T cells were pelleted by centrifugation and washed once in PBS. The pellet was then resuspended thoroughly in 1 mL of lysis buffer (20 mM Tris-HCl, pH 7.4, 150 mM NaCl, 5 mM MgCl2, 1 mM DTT, 100 μg/mL cycloheximide, 1 % v/v Triton X-100, 25 U/mL Turbo DNase I (Invitrogen)) and digested on ice for 30 min. The cell lysate was triturated through a 26 G needle 10 times and clarified by centrifugation (10 min, 4°C, 20,000 xg). Lysates were aliquoted, snap-frozen in liquid nitrogen, and stored at -80°C until ready for use.

RNA concentration of the lysates was determined using the QuantIt RiboGreen RNA kit (Invitrogen). Lysates were subsequently digested by P1 nuclease (New England Biolabs) at 20 U per µg of RNA for 1 hour at 37°C on a tube rotator. Ribosomes were pelleted by sucrose cushion in a TLA 110 rotor at 4°C for 1 hour at 100,000 rpm. Trizol was added to the ribosome pellets and RNA was extracted using a modified method called mirRICH to preferentially enrich for small RNA fragments (Choi et al., 2018; Ferguson et al., 2023). Library generation was carried out using the OTTR cDNA library construction kit (Karnateq and a generous gift from the Collins lab) following the manufacturer’s protocol. 40 ng of RNA per sample was used as input for the OTTR reaction. In parallel, 30 and 40 nucleotide control RNA oligos (a gift from the Collins lab) were used as input for the OTTR reaction to use as size selection markers (Ferguson et al., 2023). cDNA cleanup was performed following RNase A and RNase H digestion using an Oligo Clean & Concentrator kit (Zymo Research). cDNAs were size selected on a 0.6x TBE 8% Urea-PAGE and visualized using the fluorescence of the IR800 dye included in the 5’ adapter. The region between the 30 and 40 nucleotide controls was excised and the cDNA was eluted. Following the protocol described in McGlincy et al., cDNA concentrations were measured by qPCR to determine the number of PCR cycles necessary to amplify each library using Illumina primers (McGlincy & Ingolia, 2017).

Libraries were quantified using the NEBNext Library Quant Kit, pooled at equimolar concentration, and sequenced by Novogene on a NovaSeq X Plus 150PE platform. Ribosome profiling computational analysis was carried out as described by Ferguson et al., with minor modifications (Ferguson et al., 2023). First, the adapters were trimmed, unique molecular identifiers (UMIs) were annotated, and reads shorter than 20 nucleotides or of poor quality were discarded using Cutadapt:

-m 2 -a AGATCGGAAGAGCACACGTCTGAACTCCAGTCAC -A AGATCGGAAGAGCGTCGTGTAGGGAAAGAGTGT -u 7 -U -7 --rename=’{id}_NTA={r1.cut_prefix}’ -m 20 -q 10

Of note, although the libraries were sequenced with paired-end sequencing, we only used Read 1 for our analysis after this point. Reads were sequentially mapped to rRNA, ncRNA, mtDNA, mRNA, gDNA, size-selection oligos, and OTTR adapter references using bowtie2. For mRNA, reads were aligned to a custom transcriptome that includes both the NCBI RefSeq MANE v0.95 transcript set and the 13 mitochondrial mRNA sequences from Ensembl (Ferguson et al., 2023). We proceeded with further analysis using only the reads that mapped to mRNA. PCR duplicates were collapsed using UMI-tools (Smith et al., 2017). Reads mapping outside of the coding sequence (+15^th^ to -10^th^ codon) were removed using Bedtools intersect (Quinlan, 2014). The remaining reads were counted using Salmon (Patro et al., 2017). Differential translation efficiency analysis was carried out using DeltaTE (Chothani et al., 2019).

## Data Availability

The RNA-Seq and ribosome profiling datasets from this work have been deposited in NCBI’s Gene Expression Omnibus and can be found at GSE313691 and GSE313692, respectively.

## Acknowledgments

We would like to thank Dr. Nikolay Aleksashin and Cynthia Hermosillo for reviewing the manuscript and providing helpful comments. We would also like to thank the Collins lab and Dr. Lucas Ferguson in particular for sharing OTTR library generation reagents. Lastly, we’d like to thank Dr. Nicholas Ingolia for his advice on the ribosome profiling analysis and interpretation.

## Supplementary Tables and Figures

**Supplementary Table 1. Differential translation efficiency of chronic vs. reactivated T cells.** Tabs include analysis of ribosome protected fragments (deltaRPF), transcription profiling (deltaRNA), and translation efficiency (deltaTE).

**Supplementary Table 2. Differential translation efficiency of reactivated vs. acutely activated T cells.** Tabs include analysis of ribosome protected fragments (deltaRPF), transcription profiling (deltaRNA), and translation efficiency (deltaTE).

**Supplementary Table 3. Categories of transcriptional and translational changes in activated T cells**. Tabs for chronic vs. reactivated T cells and reactivated vs. acute T cells are included.

**Supplementary Figure 1.**
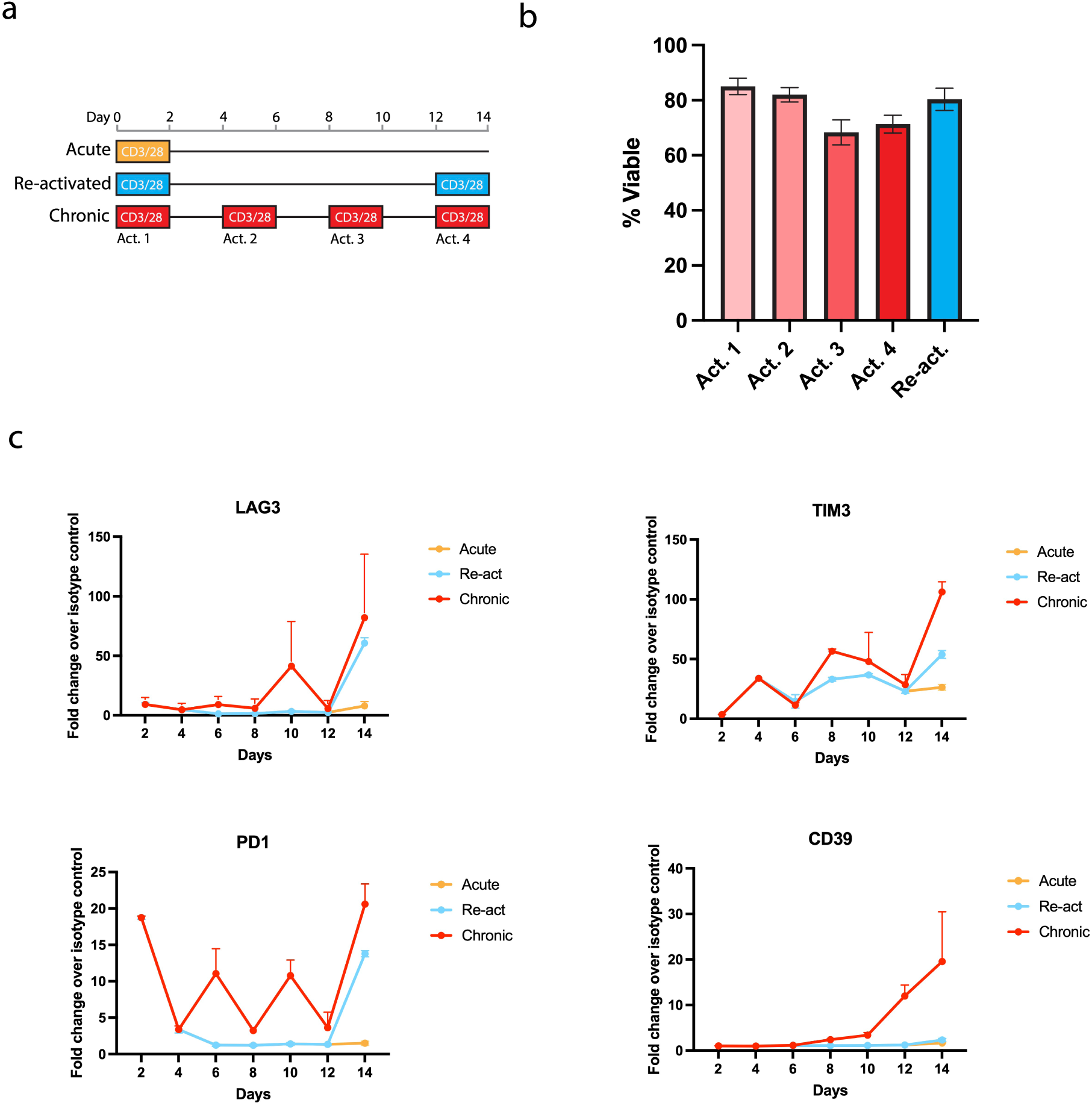
*In vitro* chronically activated CD8+ T cells show hallmarks of T cell exhaustion. **(a)** Schematic of T cell activation protocol. **(b)** Viabilities of Tchron and Tre-act cells at the end of each 2-day activation. **(c)** LAG3, TIM3, PD1, and CD39 expression measured every two days by flow cytometry.

**Supplementary Figure 2.**
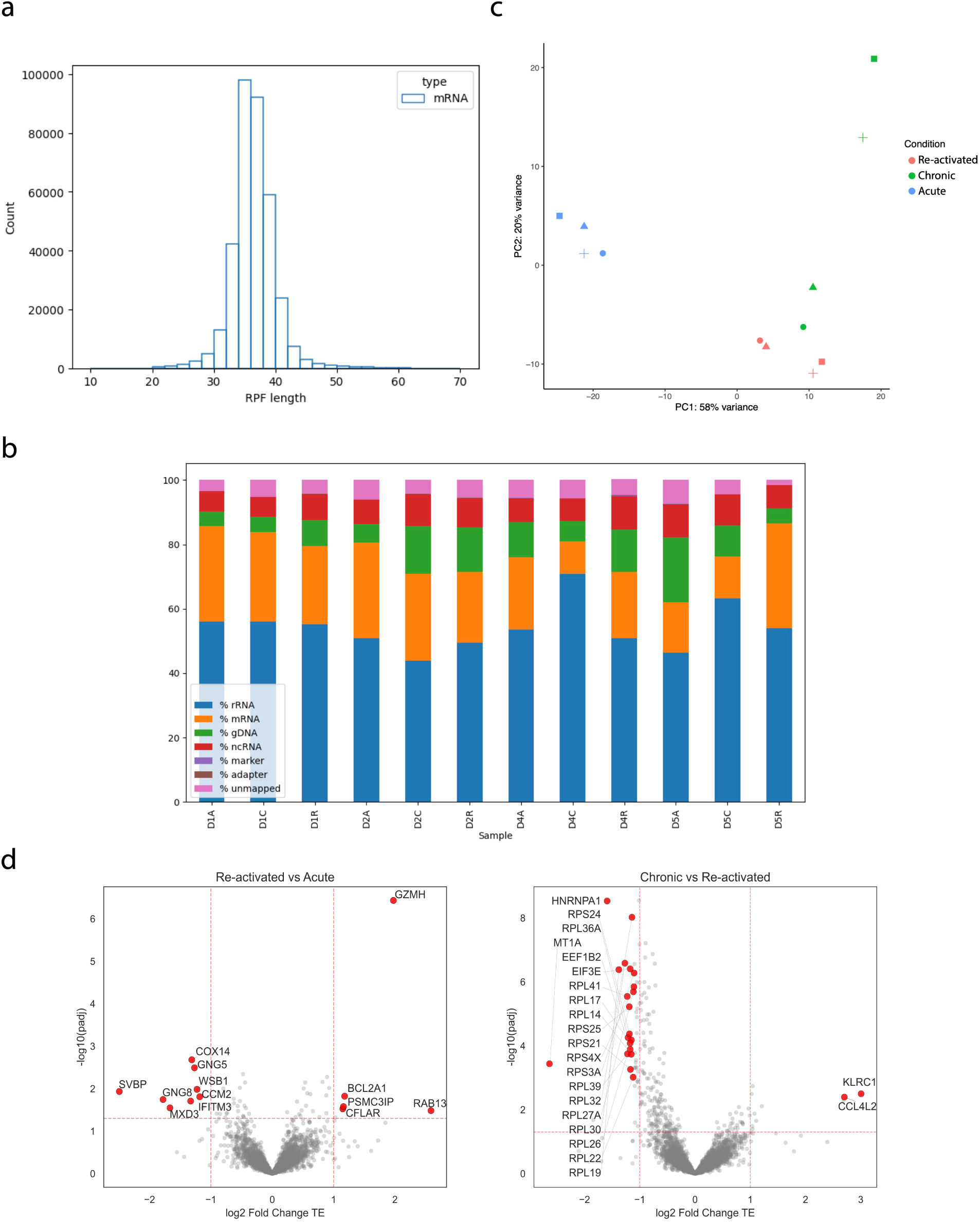
Transcription profiling and ribosome profiling statistics and comparisons. **(a)** Read length distribution of RPFs mapping to mRNA. Data represents only one donor and condition (donor 1, acute); the other donors and conditions showed similar distributions. **(b)** Percentage of RPFs mapping to various transcript types (rRNA, mRNA, gDNA, etc). **(c)** Principal component analysis of RPFs from Tacute, Tre-act, and Tchron cells in four donors. **(d)** Volcano plots of log2(fold change) TE comparing Tre-act to Tacute (left) and Tchron to Tre-act (right).

**Supplementary Figure 3.**
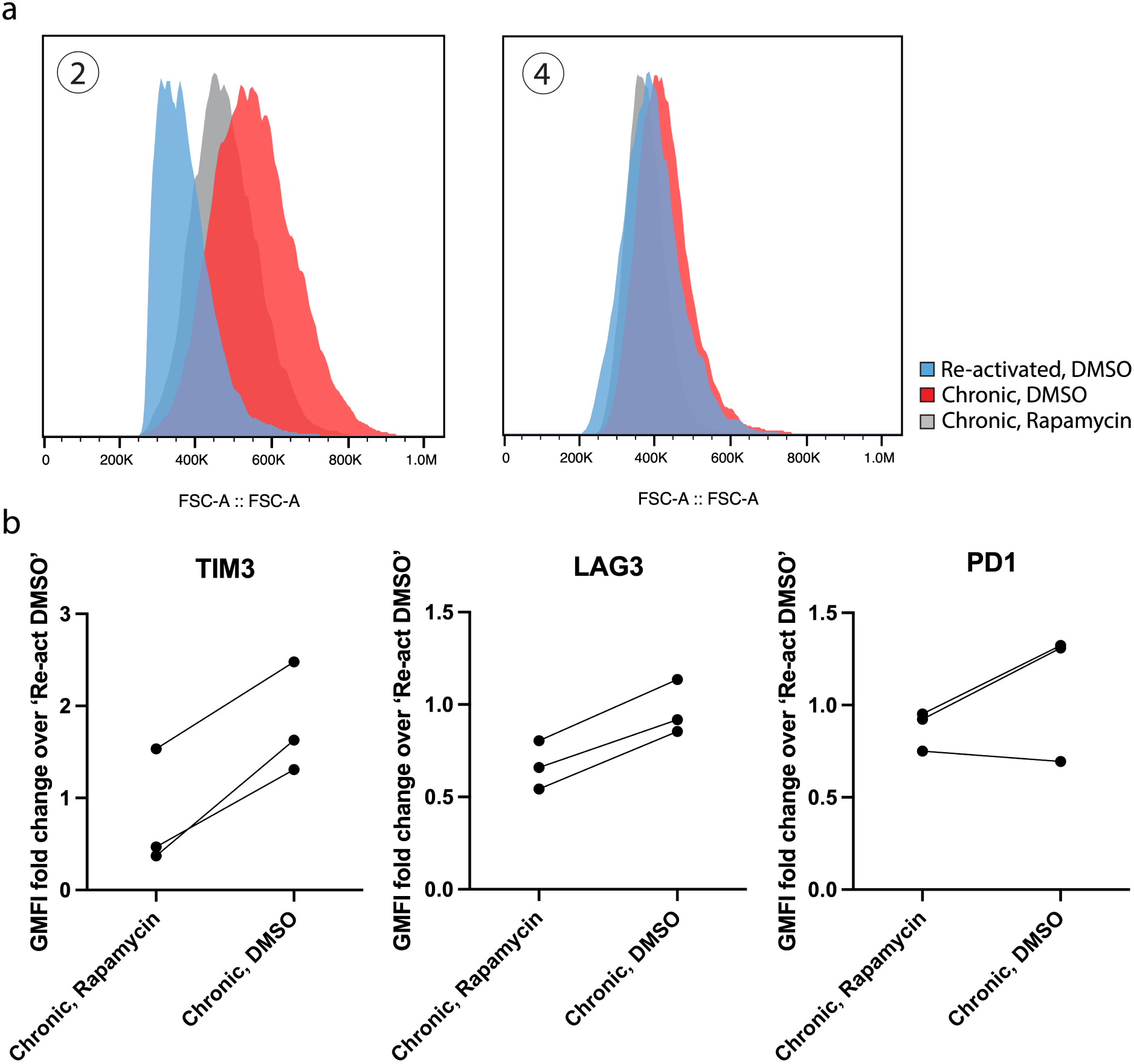
Rapamycin treatment slows the progression of T cell exhaustion. (a) Representative flow cytometry histograms of forward scatter (FSC) (n = 3 donors) at timepoints 2 (left) and 4 (right) (timepoints refer to those indicated in Figure 4a). (b) Fold change GMFI of TIM3, LAG3, and PD1 expression in Tchron cells over Tre-act cells treated with DMSO (n = 3 donors). The data shows the last timepoint (day 14).

